# Genome assemblies across the diverse evolutionary spectrum of *Leishmania* protozoan parasites

**DOI:** 10.1101/2021.05.29.446295

**Authors:** Wesley C. Warren, Natalia S. Akopyants, Deborah E. Dobson, Christiane Hertz-Fowler, Lon-Fye Lye, Peter J. Myler, Gowthaman Ramasamy, Achchuthan Shanmugasundram, Fatima Silva-Franco, Sascha Steinbiss, Chad Tomlinson, Richard K. Wilson, Stephen M. Beverley

## Abstract

We report high-quality draft assemblies and gene annotation for 13 species and/or strains of the protozoan parasite genera *Leishmania, Endotrypanum* and *Crithidia*, which span the phylogenetic diversity of the Leishmaniinae sub-family within the kinetoplastid order of the phylum Euglenazoa. These resources will support studies on the origins of parasitism.

---

*Leishmania* species are widespread parasites of mammals transmitted by biting insects. Over 1.7 billion people worldwide are at risk, with hundreds of millions of people infected (1-4). The genus comprises more than 50 species, which are primarily zoonotic but in humans cause disease ranging from mild cutaneous lesions to more disseminated forms, to fatal visceralizing disease (5). While parasitism by *Leishmania* has been intensively studied, the species-specific factors that enable mammalian or insect host infections are less well understood. To provide a broad phylogenetic snapshot, we selected a spectrum of species and strains across the Leishmaniinae sub-family (5), targeting lineages within the subgenera *Leishmania, Viannia*, and *Mundinia*, as well as allied *Endotrypanum* and the outgroup *Crithidia fasciculata* (Table 1).

**Table 1.**
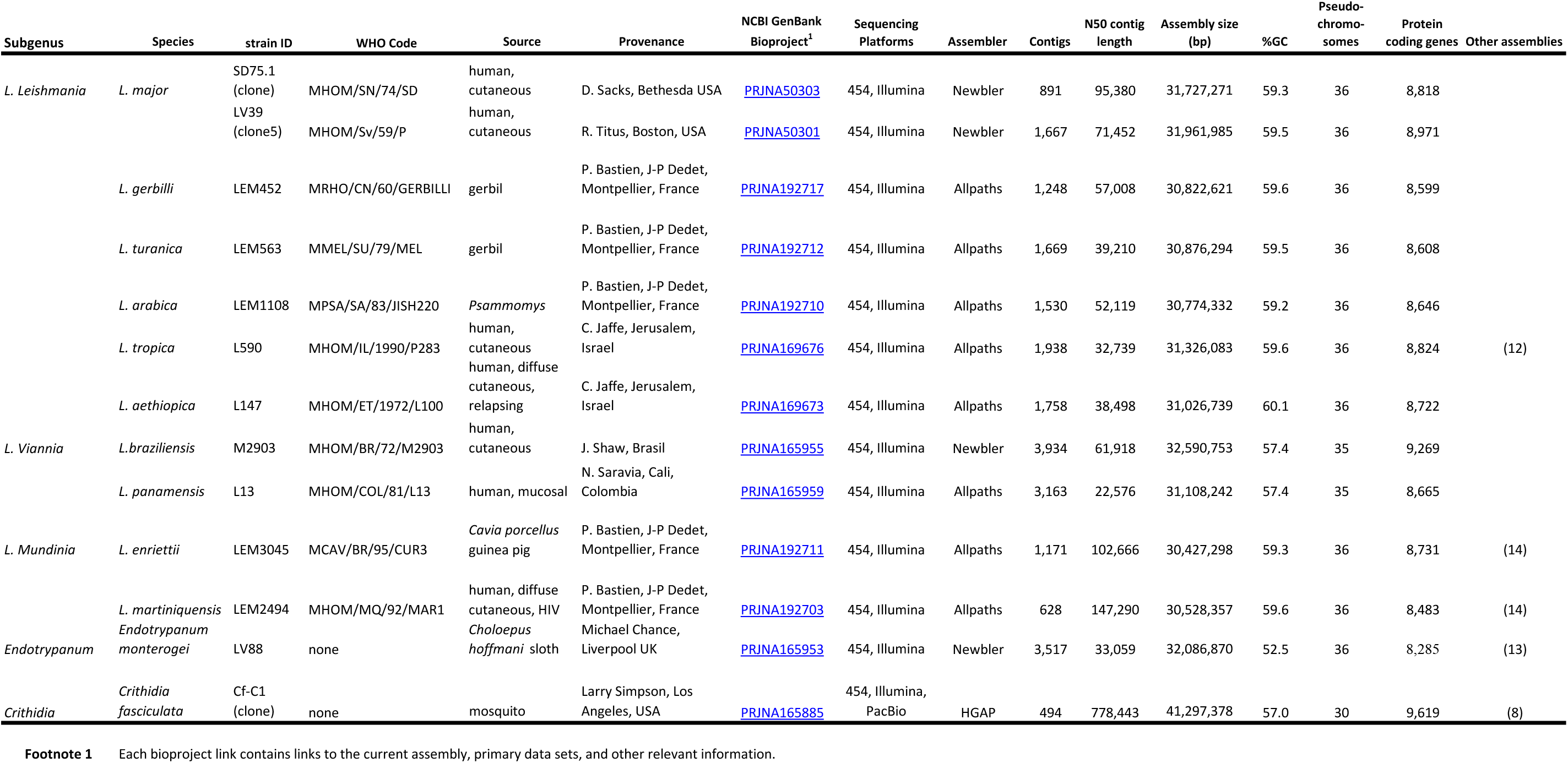
Description of *Leishmania* species and strains including assembly parameters and links.

Parasites were cultivated in M199 or Schneider’s medium (6) and grown to late log phase, before harvesting, lysis, and DNA purification by phenol/chloroform extraction and/or banding in CsCl gradients (to remove mitochondrial maxi- or minicircle DNA). Sequencing libraries were generated using the Illumina paired-end DNA sample preparation kit (PE-102-1001) according to the manufacturer’s directions. Fragment libraries of 3 and 8kb were prepared using protocols for 454 sequencing (Roche Life Sciences). Sequencing was performed on either a 454 GS FLX Titanium (average read length of 305bp; Roche 454 Life Sciences), or Illumina GAIIx and HiSeq2000 instruments (paired end 100bp read length), except for *Crithidia* which additionally utilized long-reads generated on a RSII instrument (P5/C3 chemistry; Pacific BioSciences) (7). Total sequence genome coverage on the Illumina GAIIX or HiSeq 2000 instrument was on average 105x with tiered library insert sizes (50x fragments, 45x 3kb, and 10x 8kb and 0.05x 40kb). For all Illumina sequences we used the read processing steps within the ALLPATHS-LG (8) software prior to *de novo* assembly, which incorporates read error correction methods described by Pevzner *et al* (9). Genome assemblies were performed with default parameters using Newbler v2.0.1 (10) for 454 reads, ALLPATHS-LG (8) for Illumina reads, or HGAP version 3 (11) for long-reads (Table 1). Contigs and scaffolds were organized into pseudochromosomes using ABACAS2 (https://github.com/satta/ABACAS2), a successor to ABACAS (12), by alignment to the *Leishmania major* Friedlin genome, with the exception of *L. braziliensis* M2903 and *L. panamensis*, which were aligned to the *L. braziliensis* M2904 genome. The estimated haploid genome sizes ranged from 30.4 to 41.3 megabases (13).

Gene annotations were performed using the comprehensive Companion tool which incorporates a variety of *de novo* prediction criteria, as well as information from closely related genomes when available (14). The number of protein-coding genes predicted ranged from 8,285 to 9,619, typical of other *Leishmania* (13). Full annotations, as well as a variety of tools for the visualization or analysis of these genomes, are available from TriTrypDB (www.tritrypdb.org).

## Public availability and accession number(s)

Assemblies are deposited in the NCBI GenBank Repository under the Bioproject numbers in Table 1, including links to the primary data and annotations (PRJNA50303, PRJNA50301, PRJNA192717, PRJNA192712, PRJNA192710, PRJNA169676, PRJNA169673, PRJNA165955, PRJNA165959, PRJNA192711, PRJNA192703, PRJNA165953, PRJNA165885). Chromosome builds are available through the TriTrypDB portal (http://tritrypdb.org/tritrypdb).

## Acknowledgements

NIH grant AI29646 to Stephen M. Beverley provided support for NSA, DED and L-FL, NIH-NHGRI grant HG00307907 to Richard K. Wilson supported WCW, and CT, and AI103858 to Peter J Myler supported GA. Funding to Christiane Hertz-Fowler from Wellcome Trust grants WT099198MA and WT108443MA provided support for FS-F, AS and SS. We thank Patrick Minx for assembly curation. We thank the colleagues listed in Table 1 for generously providing strains and Brian Brunk, David Roos, Thomas Otto and Matt Berriman for discussions. The WU Institutional Biosafety Committee reviewed and approved the parasite work reported here (01-015).

## Notes

### Competing Interest Statement

The authors have declared no competing interest.

### Summary of Updates

minor edits and revisions - this is the version accepted at Microbiology Resource Announcements (2021)

